# Implementation of a practical and effective control program for *Taenia solium* in the Banke District of Nepal

**DOI:** 10.1101/416370

**Authors:** Ishab Poudel, Keshav Sah, Suyog Subedi, Dinesh Kumar Singh, Peetambar Kushwaha, Angela Colston, Charles G. Gauci, Meritxell Donadeu, Marshall W. Lightowlers

## Abstract

*Taenia solium* is a zoonotic cestode parasite which causes human neurocysticercosis. Pigs transmit the parasite by acting as the intermediate host. An intervention was implemented in pigs to control transmission of *T. solium* in Dalit communities of Banke District, Nepal. Every 3 months, pigs were vaccinated with the TSOL18 recombinant vaccine (Cysvax™, IIL, India)) and, at the same time, given an oral treatment with 30mg/kg oxfendazole (Paranthic 10%™, MCI, Morocco). The prevalence of porcine cysticercosis was determined in both an intervention area as well as a similar no intervention control area, among randomly selected, slaughter-age pigs. Post mortem assessments were undertaken both at the start and at the end of the intervention. Participants conducting the post mortem assessments were blinded as to the source of the animals being assessed. At the start of the intervention the prevalence of porcine cysticercosis was 23.6% and 34.5% in the control and intervention areas, respectively. Following the intervention, the prevalence of cysticercosis in pigs from the control area was 16.7% (no significant change), whereas no infection was detected after complete slicing of all muscle tissue and brain in animals from the intervention area (P=0.004). These findings are discussed in relation to the feasibility and sustainability of *T. solium* control. The 3-monthly vaccination and drug treatment intervention in pigs used here is suggested as an effective and practical method for reducing *T. solium* transmission, thereby reducing the incidence of human neurocysticercosis.

**Author summary:** Neurocysticercosis is a disease caused by a parasitic infection of the brain. The parasite responsible, *Taenia solium*, is transmitted by pigs where human sanitation is poor and pigs roam freely. Neurocysticercosis is responsible for many cases of epilepsy in people living in poor, developing countries. The feasibility and sustainability of implementing control measures have been major impediments to reducing the incidence of neurocysticercosis. Recently, two new commercial products have become available for pigs which together offer the possibility of interrupting the parasite’s transmission the TSOL18 vaccine (Cysvax™, IIL, India) and an oxfendazole formulation (Paranthic 10%™, MCI, Morocco) licensed for use in pigs for the treatment of cysticercosis. Here we describe the impact of implementing vaccination plus drug treatment of pigs in the Banke district of Nepal. The intervention eliminated the risk of transmission of *T. solium* by the animals vaccinated and treated during the trial. Application of the vaccination and drug treatment program used here, possibly with strategic use of anthelmintics also in the human population, is an effective option for reducing the incidence of neurocysticercosis in Nepal and elsewhere.

## Introduction

Neurocysticercosis is a serious medical condition caused by infection in the brain or other nervous tissue with the larval stage of the parasite *Taenia solium*. The life cycle of the parasite involves pigs and humans in a prey-predator cycle. Humans are the obligatory definitive host for *T. solium* and harbour the adult tapeworm in the small intestine. Tapeworm eggs are released with the faeces and, if they are ingested by pigs, the larval cysticercus stage develops, principally in the muscle tissues. The life cycle is completed when humans eat insufficiently cooked, infected pig meat, leading to the development of a tapeworm. The serious medical consequences of *T. solium* infection arise because the eggs released by a tapeworm carrier are not only infective for pigs but can also cause cysticercosis if accidentally ingested by humans. In humans the cysticercus larvae have a propensity to encyst in the brain, causing neurocysticercosis, a common symptom of which is epilepsy.

The full life cycle of *T. solium* is perpetuated where sanitation conditions are poor, pigs have access to human faeces or food contaminated with human faeces, and where pork is ingested raw or poorly cooked. Hence, the full life cycle of *T. solium* is restricted to populations living in many of the poorest countries of the world. The propensity of *T. solium* to encyst in the brain of humans is responsible for the parasite causing 29% of seizure cases in areas where *T. solium* transmission occurs [1].

Human cysticercosis is one of a small number of diseases that have been formally recognised as being capable of being eradicated [2]. Improvements in sanitation and pig rearing practices in developed countries have led to a cessation in *T. solium* transmission, however attempts to institute cysticercosis control measures in poor communities have had limited success [3]. There has not as yet been an example where control activities specifically directed towards *T. solium* have had a sustained impact on disease transmission.

Control measures for *T. solium* that have been evaluated include treatment of human taeniasis cases with niclosamide or praziquantel, improvement in sanitation and other practices through public education, vaccination and medication of pigs, and improvement in pig rearing and meat inspection practices [3]. A major limitation to achieving a long-lasting reduction in neurocysticercosis has been the sustainability of *T. solium* control activities. Lightowlers and Donadeu [4] presented a logical model for control of *T. solium* transmission using a combination of vaccination and medication in pigs. Combined use of the TSOL18 vaccine [5] and oxfendazole treatment [6] in all pigs at 3-monthly intervals was predicted to lead to a cessation of *T. solium* transmission by slaughter-age pigs within a year of initiation of the program [4].

*T. solium* neurocysticercosis is a major medical concern in Nepal where it has been determined to cause the highest burden of disease due to a parasitic infection [7]. Sah et al. [8] identified the Banke District in mid-western Nepal as having a high prevalence of porcine cysticercosis. Recently two new commercially produced and registered products manufactured according to Good Manufacturing Practice guidelines have become available for control of cysticercosis in pigs the TSOL18 vaccine (Cysvax™) produced by Indian Immunologicals Limited, India, and Paranthic 10%™ an oxfendazole formulation manufactured by MCI Santé Animale, Morocco which is specifically registered for the treatment of porcine cysticercosis. These new products provide the opportunity for an assessment of a 3-monthly pig vaccination and treatment program on *T. solium* transmission. Here we describe the impact of these *T. solium* control measures implemented in pigs in the Banke district of Nepal.

## Materials and methods

### Study design

The study was conducted in Udaypur Village Development Committee (VDC) and Hirminiya & Betahani VDC of the Banke district in Nepal (81°37’E to 81°42’E, 27°90N to 28°20’N). Two distinct areas were involved, one a non-intervention area and the other an area where a combined intervention involving TSOL18 vaccination and oxfendazole treatment of pigs was undertaken. The study was designed to be able to identify an 80% reduction in the prevalence of porcine cysticercosis in slaughter-weight pigs. Sample size calculations were undertaken using a one-sided likelihood ratio test at the 5% significance level using SAS 9.3 (SAS Institute, Cary North Carolina, USA) with the TWOSAMPLEFREQ command in the PROC POWER procedure. Assuming an initial prevalence of infection of 20%, sample sizes of 55 animals were required in each area at the start and end of the trial in order to meet the desired statistical power.

Two study areas were selected, each containing a total of at least 200 pigs. The eligibility criteria for animals that were enrolled in the study were indigenous breed pigs >8 weeks of age, not heavily pregnant and not clinically ill. In the treatment area, animals that were destined for slaughter within 3 weeks were excluded in compliance with the withholding period for the oxfendazole formulation that was used [9]. At both the start and the end of the trial, a random selection of enrolled animals was assessed for cysticercosis by necropsy examination (detailed below), as this is the only reliable, sensitive and accurate method that is available to diagnose porcine cysticercosis [10].

In the intervention area, the day of first treatment administration to pigs was defined as Day 0. Pigs were enrolled and received their first treatment administration within a target period of 15 days. Subsequent administrations of treatments to pigs, including enrolment and treatment of new pigs, occurred at intervals of three months and on each occasion were completed within a target period of 15 days. The study had a duration of 12 months and hence involved 4 interventions.

### Ethics

The study was approved by the Nepal Veterinary Council and was conducted adhering to the Council’s requirements for animal husbandry.

### Enrolment of pigs in the trial and pig management practices

For the animals that met the eligibility criteria, written informed consent was obtained from the farmers. In the intervention area 114 pig rearing households agreed to participate; three households declined to participate. In both the intervention and control areas, detailed information was recorded about farming and animal management practices. Farmers provided, or estimated, the age of every pig. In the intervention area, farmers were questioned about the origin and age of all new pigs and also about what had happened to any animal that had been present for one intervention but absent at a subsequent intervention visit. Approximately 30% of piglets died before the age of 3-4 months. Classical Swine Fever (CSF) caused significant mortality. A minority of farmers vaccinate with commercial CSF vaccine this generally being the only veterinary attention that the animals received.

### Vaccination and oxfendazole treatment

The intervention team consisted of a registered veterinary doctor, two veterinary technicians and 2-4 pig catchers. After obtaining the consent of the owner, all pigs meeting the enrolment criteria were caught and numbered tags applied to both ears. Animal weight in kilograms was estimated using a measuring tape and the formula [Girth^2^ × Length/400]/2.2. The dose for oxfendazole (3ml/10kg Paranthic 10%™, MCI Sante Animale, Morocco) was calculated according to the animal weight (30mg/kg body weight) and was applied *per os*. Concurrently, 1ml TSOL18 vaccine (150μg TSOL18 recombinant protein in mineral oil adjuvant; Cysvax™, Indian Immunologicals Limited, India) was administered intramuscularly in the left side of the neck behind the base of the ear, prior to release of the animal. A different needle was used for every vaccination. The procedures were undertaken swiftly and efficiently in order to minimize stress on the animals.

### Post mortem procedures

Animals were purchased having a weight consistent with that at which pigs are commonly sold or slaughtered in the communities. Slaughter weight was determined through advice from the farmers; the mean weight of animals that the farmers indicated were available for slaughter (and which were purchased for post mortem analyses) was 70kg, although individual animal weights ranged from 35kg to more than 175kg. Necropsy procedures undertaken on 110 slaughter-weight pigs at the start of the intervention are described by Sah et al. [8]. Similar procedures were undertaken for the post mortem analyses at the end of the trial but with some variations, as follows. All staff involved in the post mortems were blinded as to whether the animals were from the control or intervention areas. Animals from both areas were necropsied in random order. The animals were transported to a licensed commercial abattoir in Nepalgunj Municipality, Banke where they were euthanized by slaughter house staff according to normal commercial practice processes. The viscera were removed and the heart, liver, lungs, both kidneys and the full diaphragm retained in numbered containers. The organs and the two halves of the carcase, including the head, were refrigerated overnight at 4 °C, after which the carcase was skinned. The tongue, masticatory muscles (both right and left sides) and brain were removed and retained. The muscles from each side of the carcase were dissected from the bones and kept separately.

### Examination for *Taenia solium* cysts

Except in cases of very heavy infection, all the retained organs and muscles of the right hand side of the carcase were sliced by hand at intervals of approximately 3mm and examined meticulously for the presence of *T. solium* cysticerci or other lesions. During the necropsies undertaken at the end of the trial, when no cysticerci were detected in the tongue, masticatory muscles, diaphragm, brain or muscles from the right hand side of the carcase, the muscles of the left hand side of the carcase were also sliced. Cysticerci were recorded as viable when they appeared as translucent vesicles filled with transparent fluid and having a visible white scolex. Non-viable lesions were recorded separately in cases where vesicles were non-translucent, containing a dense white or yellowish fluid and having no scolex and in cases of fibrosed or calcified lesions. Suspect, non-viable lesions that were not calcified were placed into RNA-later (Sigma) and investigated by PCR analysis of a fragment of the mitochondrial 12S rDNA gene using the restriction enzymes *Dde*I and *Hinf*I or *Hpa*I, as described by Rodriguez-Hidalgo et al. [11], Devleesschauwer et al. [12] and Dermauw et al. [13]. In carcases that contained thousands of cysts, all of the heart, liver, kidneys, lungs, diaphragm, tongue, masticatory muscles and brain were sliced and counted as above. The remaining carcase musculature was weighed and representative samples from different muscle sites were selected representing approximately 1kg. This sample was weighed accurately and then sliced and counted as above and the number of cysts in the carcase muscles estimated from the total muscle weight.

### Definition of a case of confirmed porcine cysticercosis

The definition of a confirmed case of cysticercosis which was adopted by Sah et al. [8] was also used here. An animal was determined to be a confirmed case of porcine cysticercosis if one or more viable *T. solium* cysticerci was found in the muscle and or the brain, or if more than one non-viable lesion was detected in the muscles and/or brain. Animals having only non-viable lesions in organs that are not typical locations for *T. solium* (eg the liver, lungs or kidneys), and which could not be confirmed as being *T. solium* by DNA analyses, were excluded. Direct comparisons of infection prevalence at the start and end of the trial were made on the basis of infections detected in the various organs examined as well as cysts found in the right hand side of the carcase, as this was the procedure used for the post mortems undertaken at the start of the trial [8].

### Data analysis

Raw data was transcribed into pre-formatted Excel spreadsheets suitable for importation into Genstat^®^ 18th edition. Statistical analysis was undertaken to evaluate the effect of treatments on the prevalence of *T. solium* cysts at post mortem. Prevalence results were compared within and between study groups, at baseline and end of study, using a two-sample binomial test. A generalised linear model with logit link function (logistic model) for binary data was also used to confirm results and to provide standard error estimates and confidence intervals around prevalence figures (Genstat^®^ 18th edition).

## Results

### Pig populations in the study areas

At the start of the trial the total pig population in the control and intervention areas was 805 pigs. In the intervention area there were 313 pigs in total, of which 227 met the inclusion criteria.

### Interventions

In the intervention area, a total of 4 rounds of pig vaccination and oxfendazole treatments were carried out between August 2016 and May 2017. A total of 585 pigs were treated during the trial, with 209, 209, 207 and 203 receiving treatment in first, second, third and fourth interventions, respectively (Tables 1 and 2).

**Table 1.**
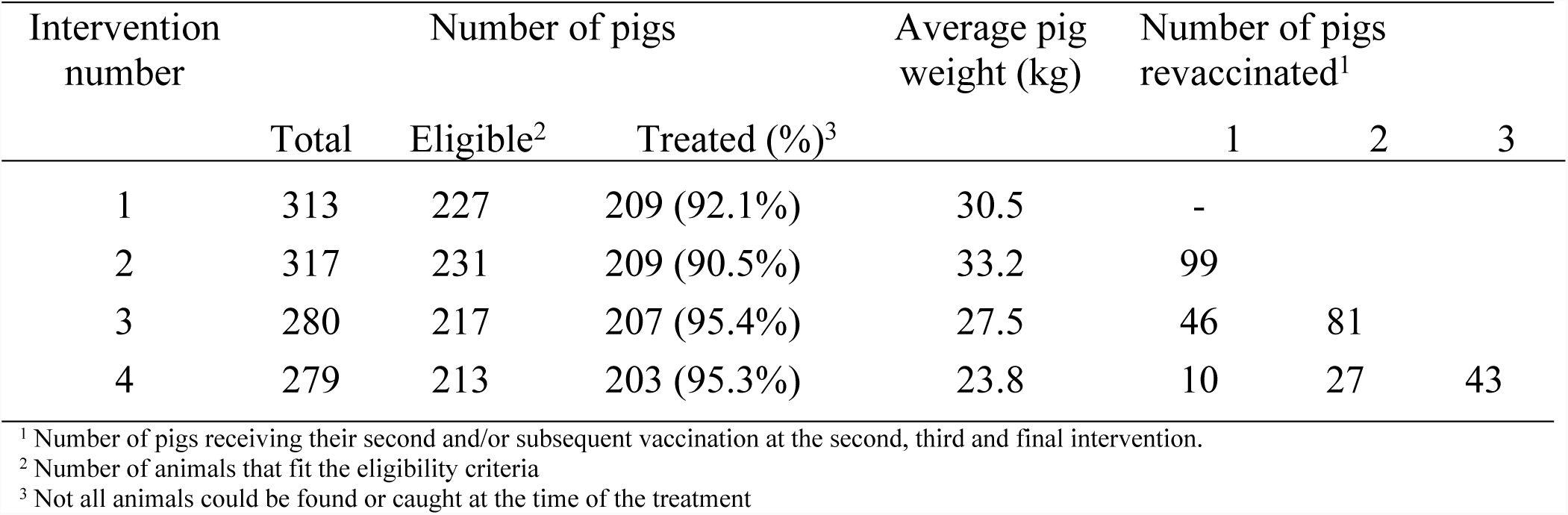
Summary of pig vaccinations and oxfendazole treatments

**Table 2.** Animals treated or not treated in the intervention area and the reasons why some animals were not treated

| Intervention | Vaccinated<br>& drug<br>treated | Pigs not treated |  |  |  | Total |
| --- | --- | --- | --- | --- | --- | --- |
|  |  | Pregnant | Sick | Age <8<br>weeks | Other<sup>1</sup> |  |
| First | 209 | 20 | 4 | 62 | 18 | 313 |
| Second | 209 | 29 | 4 | 53 | 22 | 317 |
| Third | 207 | 11 | 5 | 47 | 10 | 280 |
| Fourth | 203 | 13 | 10 | 43 | 10 | 279 |
| Total | 828 | 73 | 23 | 205 | 60 | 1189 |

<sup>1</sup> Pigs that were unable to be caught.

Interventions delivered to individual pigs are summarized in Table 3. Of the 209 animals receiving the vaccination and drug treatment during the first intervention period, 95 animals were absent at the time of the second intervention. The reasons for absence were: unable to be caught (12 animals), 2 had died, 32 had been sold or consumed locally, and 33 were otherwise ineligible for inclusion in the second round of interventions (4 were ill; 29 in late pregnancy). The remaining 16 animals were absent for unknown reasons although several were present and treated at a subsequent intervention time. Thirteen animals were not available for the second round of interventions but were present and received treatment either during the third intervention period (12 animals) or fourth intervention period (2 animals).

Three animals which were vaccinated and drug treated at the second intervention were absent at the third intervention period but were present and treated during the fourth intervention.

The number of animals which were unable to be caught decreased as the trial progressed such that approximately 95% of eligible animals received their vaccinations and oxfendazole treatments during the third and fourth rounds of intervention.

**Table 3:** Number of pigs receiving vaccination and oxfendazole treatments and the interventions in which individual animals received treatment.

| Intervention number(s) |  |  |  | No. Pigs treated |
| --- | --- | --- | --- | --- |
| 1 | - | - | - | 95 |
| 1 | 2 | - | - | 64 |
| 1 | 2 | 3 | - | 28 |
| 1 | 2 | 3 | 4 | 5 |
| 1 | - | 3 | - | 12 |
| 1 | - | 3 | 4 | 1 |
| 1 | 2 | - | 4 | 2 |
| 1 | - | - | 4 | 2 |
|  | 2 | - | - | 59 |
|  | 2 | 3 | - | 31 |
|  | 2 | 3 | 4 | 17 |
|  | 2 |  | 4 | 3 |
|  |  | 3 | - | 93 |
|  |  | 3 | 4 | 20 |
|  |  |  | 4 | 153 |
| TOTAL |  |  |  | 585 |

### Veterinary interventions, adverse reactions and farmer attitudes to the intervention

During the trial there were 28 pigs which required treatment for wounds (including 13 ear wounds, six scrotal wounds), 11 for respiratory infections and one animal noted with a neurological condition. Treatments administered were long acting oxytetracycline or sulphonamide/ trimethoprim, meloxicam as well as topical treatment of the wounds.

No adverse reactions to either vaccination or oxfendazole treatment were noted by the field staff or reported by the farmers (who were asked specifically about the issue). No injection site lesions were noted. Many farmers however were reluctant to have ear tags placed on their animals, especially since some animals developed ear infections following the first intervention. There was no reluctance on the part of the farming community to their animals being either vaccinated or given anthelmintic drench. Farmers were pleased to see worms voided in the feces after the animals were treated.

### Post mortem examination

The pigs that underwent post mortem examination at the end of the trial were from 8 to 48 months of age and weighed from 35 to 175 kilograms, mean 89kg. The number of interventions and individual treatments received by the animals in the intervention area that underwent post mortem at the end of the study, are summarized in Table 4.

**Table 4.** Records of the 33 animals from the *T. solium* intervention area that were assessed in post mortem.

| Number of interventions | Number of pigs | Intervention(s) received |  |  |  |
| --- | --- | --- | --- | --- | --- |
| 1 | 1 |  |  | 1 |  |
| 2 | 1 | 1 |  | 3 |  |
|  | 1 | 1 | 2 |  |  |
|  | 2 |  | 2 | 3 |  |
|  | 2 |  | 2 |  | 4 |
|  | 10 |  |  | 3 | 4 |
| 3 | 5 | 1 | 2 | 3 |  |
|  | 1 | 1 | 2 |  | 4 |
|  | 1 | 1 |  | 3 | 4 |
|  | 8 |  | 2 | 3 | 4 |
| 4 | 1 | 1 | 2 | 3 | 4 |

Shown are the number of interventions the individual animals received and at which intervention period(s) they were treated.

Thirty three animals from the intervention area were examined at most mortem, among which one animal had received a single intervention treatment, 16 animals had received 2 intervention treatments, 13 had received 3 treatments and 1 animal had received all 4 treatments.

### Prevalence and intensity of *T. solium* infection

The numbers of animals recorded as having *T. solium* infection in animals from the control and intervention areas, assessed at necropsy at the start and the end of the trial, are summarized in Table 5. Among the animals necropsied at the start of the trial, nineteen out of 55 (34.6%) pigs were positive from the intervention area, and thirteen out of 55 (23.6%) pigs were positive from the control area (not significant, p=0.207). The total prevalence of porcine cysticercosis (PC) was 29.1%. Approximately 9-10 months after the first intervention, 33 pigs from the intervention area and 35 pigs from the control area were subjected to post mortem to determine the presence of *T. solium* cysts. Zero out of 33 (0%) pigs were positive from the intervention group and six out of 35 (17.1%) pigs were positive from the control group (significant, p=0.004). The pre-intervention prevalence of infection was significantly greater than the post intervention prevalence in the intervention area (p<0.001). In the control area there was no significant difference between the baseline and end of study prevalence of infection with *T. solium* (p=0.424).

**Table 5.** Prevalence of *T. solium* infections detected in pigs from the intervention and control areas of Nepal

| Study area | Baseline necropsies<sup>1</sup> |  |  | End of study necropsy |  |  |
| --- | --- | --- | --- | --- | --- | --- |
|  | Total animals necropsied | Number infected | % positive | Total animals necropsied | Number infected | % positive |
|  |  |  |  | Including RHS carcase<sup>2</sup> |  |  |
| Intervention | 55 | 19 | 34.5% | 33 | 0 | 0% |
| Control | 55 | 13 | 23.6% | 35 | 6 | 17.1% |
|  |  |  |  | Including complete carcase |  |  |
| Intervention |  |  |  | 33 | 0 | 0% |
| Control |  |  |  | 35 | 8 | 22.9% |

<sup>1</sup> Data from Sah et al. (2017); <sup>2</sup> Heart, masseters, tongue, diaphragm, brain, liver, lungs plus right hand side skeletal musculature.

The number of both viable and non-viable cysts was counted according to the criteria specified for a confirmed case of *T. solium* infection, including the number in the full carcass estimated by doubling the cyst number found in the skeletal muscles of the right hand side of the carcase. The baseline post mortems revealed 8,347 (viable 7,379: non-viable 968) cysts in animals from both the control and intervention areas, with 7,039 cysts (viable 6,694: non-viable 345) in 19 pigs (average per infected animal 370.5±537.5) from the intervention area and 1308 cysts (viable 685, non-viable 623) from 13 pigs (average 100.6±145.0) from the control area. There were fewer cysts found at the end of study post mortems, with 120 cysts identified in six pigs (average 20.0±24.0) from the control area and none found to be infected in the intervention area. All animals that were recorded as having no *T. solium* infection detected in the heart, masseters, tongue, right hand carcase musculature, brain, liver or lungs had the remaining carcase musculature (the left hand side) sliced to determine whether there was any infection in the carcase at all. Two further animals were identified as being infected from the control area; one with a single viable cyst and one animal with two viable cysts only in the skeletal muscles from the left hand side of the carcase. No infection was detected, either viable or non-viable cysts, in any of the 33 animals from the intervention area after complete dissection of the carcases (Table 5).

A total of 145 pigs that had no vaccination or drug treatment (110 animals at the start of the trial plus 35 from the control region at the end of the trial) were subjected to detailed necropsy. This included careful slicing of the entire brain. Nine of these animals had one or more viable *T. solium* cysticerci in the brain. All those with cysts in the brain also had viable cysts in one or more muscle tissues. None of the 33 pigs from the intervention area had any *T. solium* cysts in the brain.

Analysis of the characteristics of all animals that were found to be infected with *T. solium* (33 at the start of the intervention and 21 at the end of the intervention; Supplementary Data) in comparison to the total number of animals that were necropsied at baseline and from the control area at the end of the trial, found no significant relationship between the age of the animals and the proportion of infected animals in the age range between 7 and >19 months of age (Pearson’s correlation coefficient 0.189, P=0.76; Fig 1). There was also no significant relationship between the percentage viability of cysts found in individual animals and the age of the pigs (Pearson’s correlation coefficient 0.188, P=0.28), nor between the intensity of infection (total number of cysts) and the age of the animal (Pearson’s correlation coefficient 0.133, P=0.45).

**Fig 1.**
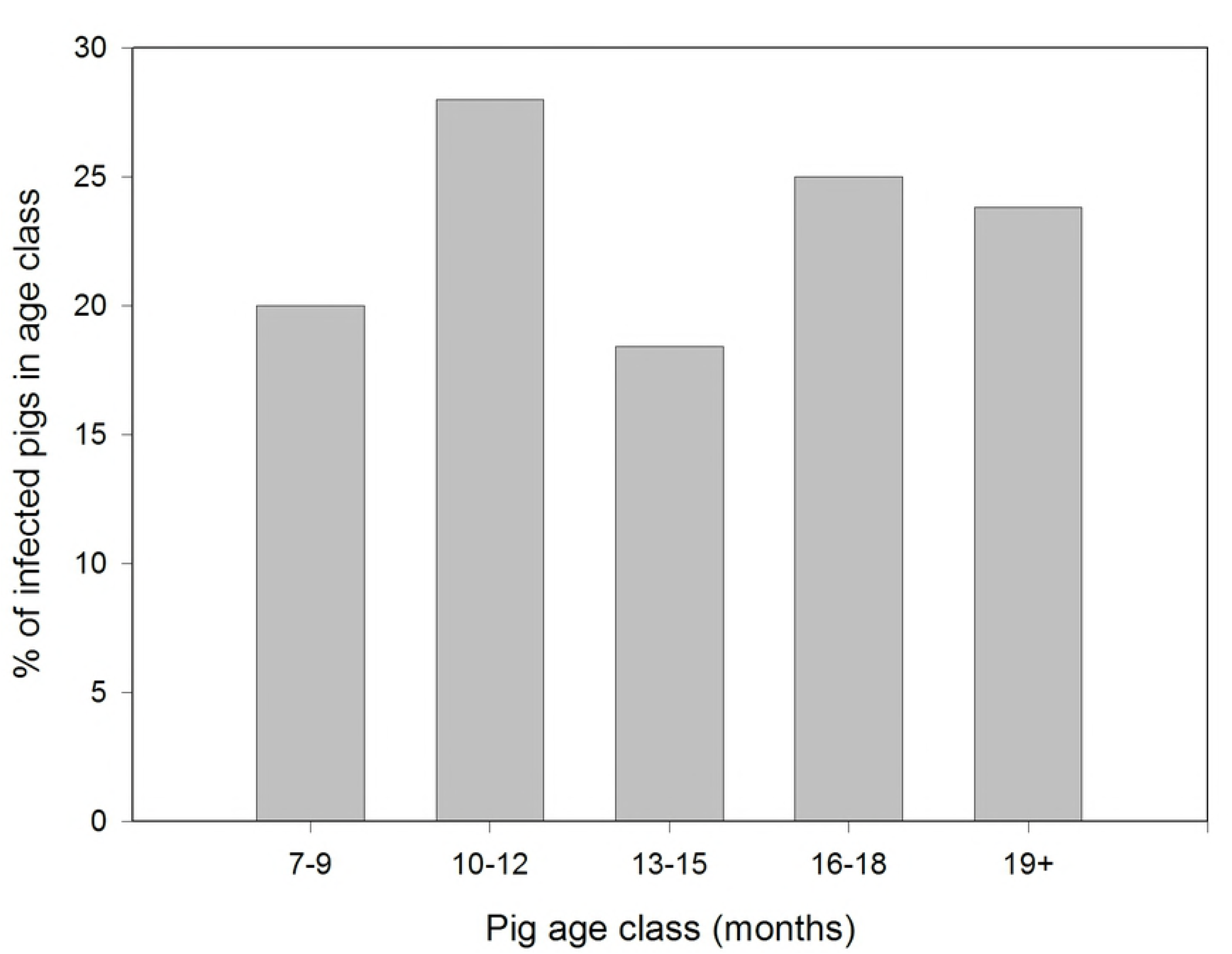
Proportion of pigs in different age classes from the Banke district of Nepal found to be infected with *Taenia solium*. Pigs of the different age classes shown were selected at random from a population of animals naturally exposed to infection in the Banke District of Nepal and assessed for infection by comprehensive investigations undertaken at necropsy.

## Discussion

Following implementation of 3-monthly treatments of the pig population in a *T. solium* endemic region of Banke District, Nepal, transmission of the parasite was eliminated among the animals that were assessed at the end of the study. Comparison of the number of infected animals found in the intervention and control areas indicates that the intervention led to a significant reduction in porcine cysticercosis (P=0.004; Table 5). This change in the risk of *T. solium* transmission is also evident when comparing the starting prevalence of infection in the intervention area with the prevalence of infection in the same area seen at the end of the intervention (P<0.001). There was a reduction in the prevalence of *T. solium* prevalence between the start and the end of the trial in the control area, however this was not significant (P=0.424). Fewer animals were able to be purchased for necropsy at the end of the trial than had been intended. Nevertheless, significant differences were seen between the prevalence of infection at the start and end of the trial, as well as between the intervention and control areas at the end of the trial, due to the higher prevalence of infection than expected at the start and the magnitude of the intervention’s impact on cysticercosis transmission.

The 3-monthly vaccination and oxfendazole treatment regime was implemented over an approximately 10 month period prior to the post mortem assessments being undertaken at the end of the trial. As the animals selected for assessment were based on them being of a size and age at which pigs from the area are generally sold or slaughtered, most of the animals assessed had been present for at least two of the interventions (Table 4). The treatment schedule which was assessed was determined to be the most effective in an area where animals are consumed from 7 months of age [14]. Available evidence suggests that immunity stimulated by the TSOL18 vaccine requires two immunizations with the currently available vaccine [15]. Excellent secondary responses to the vaccine are seen when the interval between vaccinations is between one and four months [16], hence the vaccination scheme adopted in this trial would be predicted to provide a high level of immunity. The level of protection achieved in this trial is similar to what was achieved in a previous field trial undertaken in Cameroon which involved a cohort of animals, rather than an on-going intervention program that was implemented in this trial in Nepal [17].

An estimate of the age at which animals were sent for slaughter was determined from the information obtained about animals that were present for the second intervention but had been sold prior to the third intervention, together with those that were present at the time of the third intervention but had been sold prior to the fourth intervention. Excluding animals that were absent for reasons such as them having died, being pregnant or ill, this information provided the age of the animals at which they were sold. On this basis, the typical age at slaughter of pigs in these communities was found to be 14 months when the animals were at least 60kg.

Data obtained from the animals that were not part of the intervention provide valuable information about the age at which pigs acquire *T. solium* infection in a natural endemic situation. Very little information is available concerning this topic. No significant relationship was evident among all the animals that were confirmed to have *T. solium* infection between the age of the animals and the proportion that was infected (Fig 1). Assuming that the infections acquired in young pigs persist, these data suggest that pigs acquire infection relatively early in life and that additional infections do not accumulate as the animals age. An hypothesis that would be consistent with these data would be that pigs older than approximately 1 year are relatively resistant to infection. Age-related resistance to infection is recognised in the intermediate hosts of other *Taenia* species [18].

None of the animals from the intervention area in Nepal were found to have *T. solium* in the brain tissue, whereas 9 of 145 untreated pigs were found to have cysts in the brain. This difference is not statistically significant, however the absence of cysts in vaccinated and treated pigs is consistent with the results of previous trials with the TSOL18 vaccine in which vaccinated animals also had no cysts in the brain [5, 17, 19].

The post mortem investigations undertaken at the start of the trial in Nepal included slicing of half the carcase musculature in addition to the heart, masseters, diaphragm, tongue, brain, liver, kidneys and lungs. Post mortems undertaken at the end of the trial involved the same procedures, such that direct comparisons could be made between results from the two sets of data. However, for all animals in which no cysticerci were found during the post mortems carried out at the end of the trial, the remaining musculature (left hand side carcase) was also sliced. In the case of the animals from the intervention area, none was recorded as having any cysts in the entire carcase musculature or other tissues that were examined. In the 36 control animals, two additional infected animals were identified, one having a single viable cyst and the other having two viable cysts in the left hand side musculature, but no cysts elsewhere. In this group of 36 infected animals from the control area most had light infections. Identification of infected animals by slicing muscles only from one side of the carcase, rather than the entire carcase musculature, would have missed 25% of the infected animals (2 of 8 infected). Chembensofu et al. [20] found that slicing predilection sites plus only one side of the carcase musculature would have missed 16% of the infected animals in their study undertaken with naturally infected pigs in Zambia.

During the post mortems all carcase lesions and lesions in the brain, liver, kidneys and lungs were examined for the presence of a cysticercus. No cysticerci were found other than in striated muscle tissue and the brain. Necrotic lesions and other suspect lesions were investigated for the presence of taeniid or *T. solium* DNA. No *T. solium* lesions were identified by these methods other than in the brain and striated muscle. These data are consistent with the tissue distribution of *T. solium* cysticerci in many previous studies, including the comprehensive investigations undertaken by Boa et al. [21] on naturally infected pigs in Tanzania, the majority of which were heavily infected. These data, however, contrast with those reported by Chembensofu et al. [20], who found large numbers of viable, DNA-confirmed cysticerci in the liver, lungs and kidneys of many pigs from Zambia. There is no clear explanation for this discrepancy.

Potentially effective control measures for *T. solium* have been available now for decades and yet they have not been implemented anywhere as specific strategies that have led to a sustained reduction in neurocysticercosis [3]. Feasibility and sustainability of control measures have been the stumbling block to controlling *T. solium*. The requirement for a 21-day withholding period after oxfendazole treatment of pigs, creation of necrotic lesions in the meat of drug-treated, infected pigs, and difficulties with reliably predicting the time of sale or slaughter, prevent a treat-immediately-before-slaughter approach being used in pigs to control *T. solium*. To be effective, the frequency with which intervention would need to be undertaken in the pig population is governed by the rapid turnover of pigs in the communities and constant introduction of new, susceptible animals into the population due to pigs breeding throughout the year.

Combining both vaccination and oxfendazole as a preventative treatment for porcine cysticercosis has several advantages. Firstly, the drug treatment eliminates any viable cysts that may be in an animal’s musculature prior to the animal being protected after vaccination. Secondly, the drug treats many nematode and trematode infections, as well as cysticercosis, likely providing a health and productivity boost to the treated animals [22, 23]. Oxfendazole treatment does not provide any protection for uninfected pigs against subsequent exposure to the *T. solium*, hence combined use with the vaccine provides both treatment as well as prevention from subsequent infection. After treatment of an infected pigs with oxfendazole necrotic lesions are evident in the musculature for a period of at least several weeks; the great majority disappearing within a period of 3 months [6, 24, 25]. It seems likely that some of the animals from the intervention area in Nepal that underwent post mortem investigation would have been infected with *T. solium* prior to them being fully vaccinated. However no non-viable lesions were detected in the muscles of the animals that had participated in the interventions. The three-monthly treatment regime that was implemented in the trial appears to have allowed sufficient time for any lesions that were the result of the death of parasites in muscles after medication to be resorbed before the animals reached slaughter age.

A limitation to the use of oxfendazole as a treatment for porcine cysticercosis is the requirement for a 3 week withholding period after treatment before slaughter due to the presence of drug residues in the tissues [9]. In the intervention described here, all animals ≥2 months of age were treated (other than sows near parturition). Farmers were requested to not sell or kill the treated animals for 3 weeks after each treatment. This imposes a significant burden on the farmers, especially when the procedure is repeated every 3 months, and it is difficult to monitor compliance. Also, the farmer’s requirements about selling animals can change rapidly; a family illness or other unforeseen event can impose an urgent need to sell animals. The *T. solium* transmission modelling presented by Lightowlers and Donadeu [4] predicted that a 3-monthly program of vaccination plus oxfendazole treatment of pigs between one and 7 months of age would eliminate *T. solium* infection entirely from pigs >7 months of age such that they would not require further oxfendazole treatment. Cessation of drug treatment of animals that are approaching slaughter age would reduce or prevent the risk that animals with high levels of drug residues could be consumed as well as reducing the cost. The intervention program that was applied in the trial in Nepal involved animals of all ages (>2 months). Introduction of a new 3-monthly treatment program in an area would necessarily involve all pigs to start, so as to treat and protect all the existing animals. However, immunity induced by 2 immunizations with the Cysvax vaccine, together with a natural resistance to *T. solium* infection in animals >1 year of age (mentioned above), may allow a continuing program involving vaccination plus oxfendazole medication to be effective if it were only implemented in animals up to 7 months of age [4].

The intervention that was undertaken in this trial in Nepal was relatively simple. Groups of 5-6 persons travelled by motorbike. The most time-consuming aspect of the intervention was catching the animals; having caught a pig, vaccination and drug treatment took just a few moments. The older animals were generally the more difficult and time consuming to catch. Based on the experience gained in conducting this trial in Nepal, teams of 5-6 persons who were undertaking a similar, but on-going cysticercosis control program implemented in pigs up to 7 months of age, would be able to vaccinate and drug treat approximately 100 pigs per day in Dalit communities such as those in the Banke district.

The three-monthly intervention scheme adopted here was predicted to, and did, lead to the cessation of the risk of *T. solium* transmission by the vaccinated and drug-treated animals. Any intervention limited to the pig population would not immediately affect the incidence of cysticercosis in humans because it would take time for the prevalence of human *T. solium* taeniasis to decline as new cases of taeniasis were prevented due to the absence of cysticercosis in pigs. Calculations based on the rate of re-establishment of taeniasis following mass treatment of communities [26] suggest that *T. solium* tapeworms have a lifespan of 2-3 years; a lifespan of less than 5 years is also suggested by epidemiological evidence [27]. If this were the case, implementation of an on-going intervention only in pigs would lead to a substantial reduction in, or elimination of, the incidence of human cysticercosis within about 2-3 years. Alternatively, a single treatment of the human population for taeniasis after porcine cysticercosis was controlled, would lead to a more immediate reduction in the incidence of human cysticercosis [3, 4]. Although it was not tested here, evidence about the duration of protection afforded by the TSOL18 vaccine [17] and *T. solium* transmission modelling [4] would suggest that a 3-monthly vaccination and drug treatment regime would be effective if applied only to animals up to 7 months of age, with re-vaccination only of animals kept for long periods, for example, for breeding purposes. We propose that this would be an effective, relatively simple and feasible control strategy for *T. solium* which could be applied to reduce the incidence of neurocysticercosis in Nepal and elsewhere. The feasibility of this approach has been enhanced by the availability of an effective vaccine and medication, with both becoming available, for the first time, as commercial products licensed for use in pigs for *T. solium* cysticercosis.

## Acknowledgements

The authors acknowledge the logistic supports made available by the Heifer Nepal team and administrative support from District Livestock Service Office (DLSO), Nepalgunj, Banke. Valuable contributions were made by the following government veterinary officers: Dr Bed Bahadur KC Senior Veterinary Officer, District Livestock Service Office, Banke, Nepal, Dr Krishna Raj Pandey Senior Veterinary Officer, Regional Veterinary Laboratory, Surkhet, Nepal, Dr Sankar Pandey Senior Veterinary Officer, Animal Quarantine Office, Nepalgunj, Nepal and Dr Dinesh Kumar Yadav Animal Scientist, National Agriculture Research Council, Khajura, Nepal. Participants in the necropsy team included Dr. Nirjal Dhakal, Dr. Deepak Lamsal, Dr. Sauroj Shrestha, Bishal Chand, Rakesh Chand, Niraj Dhakal, Manish Gautum, Bedika Ghising, Santosh Gyawali, Rupak Kandel, Romi Kunwar, Jitendra Lama, Bishal Maharjan, Sikesh Manandhar, Sanjay Poudel, Saurav Shrestha and Kaberi Sing, Richa Singh and Simon Singh. We thank Ms Jane Poole for statistical analyses and Ms Lizzie Chesang for data curation. The findings and conclusions contained within are those of the authors and do not necessarily reflect positions or policies of the Bill & Melinda Gates Foundation nor the UK Government.

